# Impaired Motor Awareness of Balance Control is Associated with Postural Instability in Parkinson’s Disease

**DOI:** 10.64898/2026.04.08.716824

**Authors:** Hiroyuki Hamada, Aki Takamura, Tetsuya Hasegawa, Wen Wen, Yoshihiro Itaguchi, Ken Kikuchi, Arito Yozu, Jun Ota, Arata Nakamura, Hiroaki Fujita, Keisuke Suzuki, Atsushi Yamashita, Qi An

**Author notes:** **Correspondence to:** Hiroyuki Hamada, Department of Human and Engineered Environmental Studies, Graduate School of Frontier Sciences, The University of Tokyo, 5-1-5 Kashiwanoha, Kashiwa, Chiba 277-8563, Japan.

## Abstract

**Background:** Balance instability is a major contributor to disability and falls in people with Parkinson’s disease (PwP) and is often insufficiently explained by motor impairment alone. Altered awareness of motor control has been suggested to contribute to sensorimotor dysfunction in PwP, but its relationship with balance performance is poorly understood.

**Objective:** To determine whether awareness of balance control, assessed using a control detection task (CDT), differs between healthy controls (HC) and PwP, and whether CDT performance is associated with balance-related measures.

**Methods:** Healthy older adults (n=20) and PwP (n=22) performed a standing version of the CDT based on center-of-pressure (COP) control, using a force plate. CDT accuracy was used as the primary outcome measure. Static balance during quiet standing was assessed using the COP trajectory length and rectangular area. Dynamic standing balance was assessed using the Index of Postural Stability (IPS). Group differences were examined by independent-samples t-tests. Correlations between CDT accuracy and balance measures were analyzed.

**Results:** The PwP group showed significantly lower CDT accuracy. Higher CDT accuracy was associated with better static balance in the HC group and the combined sample, and with higher IPS primarily in the PwP group.

**Conclusions:** Motor awareness during postural tasks is altered in PwP and is associated with balance control. These findings suggest that balance instability in Parkinson’s disease may involve altered balance-related action–outcome monitoring in addition to motor dysfunction.

## Introduction

Parkinson’s disease (PD), a progressive neurodegenerative movement disorder that primarily affects the motor system, is a result of the degeneration of dopaminergic neurons in the brain’s substantia nigra pars compacta. Its clinical presentation is characterized by cardinal motor symptoms, including rest tremor, rigidity, bradykinesia, and postural instability [1]. In addition to these motor deficits, PD encompasses a broad spectrum of non-motor manifestations such as autonomic dysfunction and cognitive decline, reflecting widespread neurodegeneration involving dopaminergic and non-dopaminergic systems that include noradrenergic, serotonergic, and cholinergic pathways [2].

Among the motor symptoms observed in people with Parkinson’s disease (PwP), disturbances in balance control and freezing of gait (FOG) are particularly prominent, with prevalence rates exceeding 70% at advanced stages of PD [3,4]. Falls related to impaired balance are associated with injury, reduced mobility, and loss of independence [5–7], underscoring balance impairment as a central and functionally critical feature of PD.

Impairments in balance control and gait are often only partially responsive to dopaminergic medication and may persist or worsen with the progression of PD [8,9]. This suggests that balance dysfunction among PwP cannot be explained solely by motor execution deficits; it has been speculated that postural instability and related gait disturbances are likely to involve higher-order processes supporting the integration of motor intentions and sensory information [10].

Adaptive balance control relies on the precise regulation of the body’s center of mass through the integration of multisensory inputs, i.e., visual, vestibular, and somatosensory information, together with predictive motor processes [11,12]. During voluntary postural adjustments, the nervous system generates predictions about the expected sensory consequences of intended movements and compares them with incoming sensory feedback [13,14]. Accurate correspondence among intended actions, predicted outcomes, and actual sensory consequences is essential for maintaining postural stability and enabling adaptive balance control. It has been demonstrated that PwP exhibit impairments in sensorimotor integration relevant to balance control, including predictive mechanisms, sensory reweighting, and sensory feedback processing [15,16]. Although peripheral sensory impairments have been observed in PwP, these deficits are typically mild or subtle in most cases [17]. In contrast, accumulating evidence has revealed abnormalities in central sensory processing and multisensory integration that may impair the detection and correction of mismatches during postural control [18,19].

In parallel with these sensorimotor alterations, evidence also suggests changes in motor awareness in PwP. Altered perception of movement timing and direction has been reported, indicating disruptions in the comparisons of predicted motor outcomes with sensory feedback [20]. These findings raise the possibility that balance disturbances in PwP may involve altered monitoring of self-generated postural actions in addition to motor execution deficits.

Despite the clinical importance of balance instability and gait impairment, the relationship between balance deficits and motor awareness in PwP remains unclear. It is unknown how PwP evaluate their own balance control during voluntary postural adjustments or whether altered motor awareness contributes to functional impairments in balance and gait.

We thus conducted the present study to investigate differences in the motor awareness of balance control between healthy individuals and PwP. We assessed performance on a control detection task (CDT) [21] and examined the association between CDT performance and objective balance measures. Grounded in sensorimotor integration and predictive coding frameworks, the CDT has been widely used to examine action–outcome monitoring in motor tasks, typically seated upper-limb movements under altered visual feedback [21,22].

However, findings from these paradigms cannot be assumed to generalize directly to balance control. The motor control of whole-body posture differs from upper-limb control in its reliance on trunk coordination and multisensory integration for regulating the center of mass [23]. In addition, balance requires a continuous, gravity-dependent regulation of the body’s orientation, which may impose distinct demands on predictive and evaluative mechanisms. Given that postural instability and gait disturbances are major disabling features among PwP, it is important to directly examine performance on such tasks during balance control. In the present study, we extended the CDT to a postural context by developing a standing version based on center-of-pressure control using a force plate, which enabled the assessment of self–other attribution during whole-body balance control.

Based on research demonstrating altered action–outcome monitoring in PwP [20,22], we hypothesized that (*i*) accuracy on the CDT would be reduced in PwP compared to healthy controls, reflecting impaired action–outcome monitoring during balance control, and (*ii*) reduced CDT accuracy would be associated with poorer balance performance.

## Methods

### Participants

Twenty HC (73.6 ± 6.0 years old, 11 females, 9 males) and 22 PwP undergoing short-term hospitalization for rehabilitation (73.5 ± 7.7 years old, 12 females, 10 males) participated. Group characteristics are summarized in Table 1. This study was performed in accord with the principles of the Declaration of Helsinki, and approval for the study was granted by the Ethics Committee at The University of Tokyo (approval no. KE23-68). Each participant provided written informed consent prior to participation.

**Table 1.**
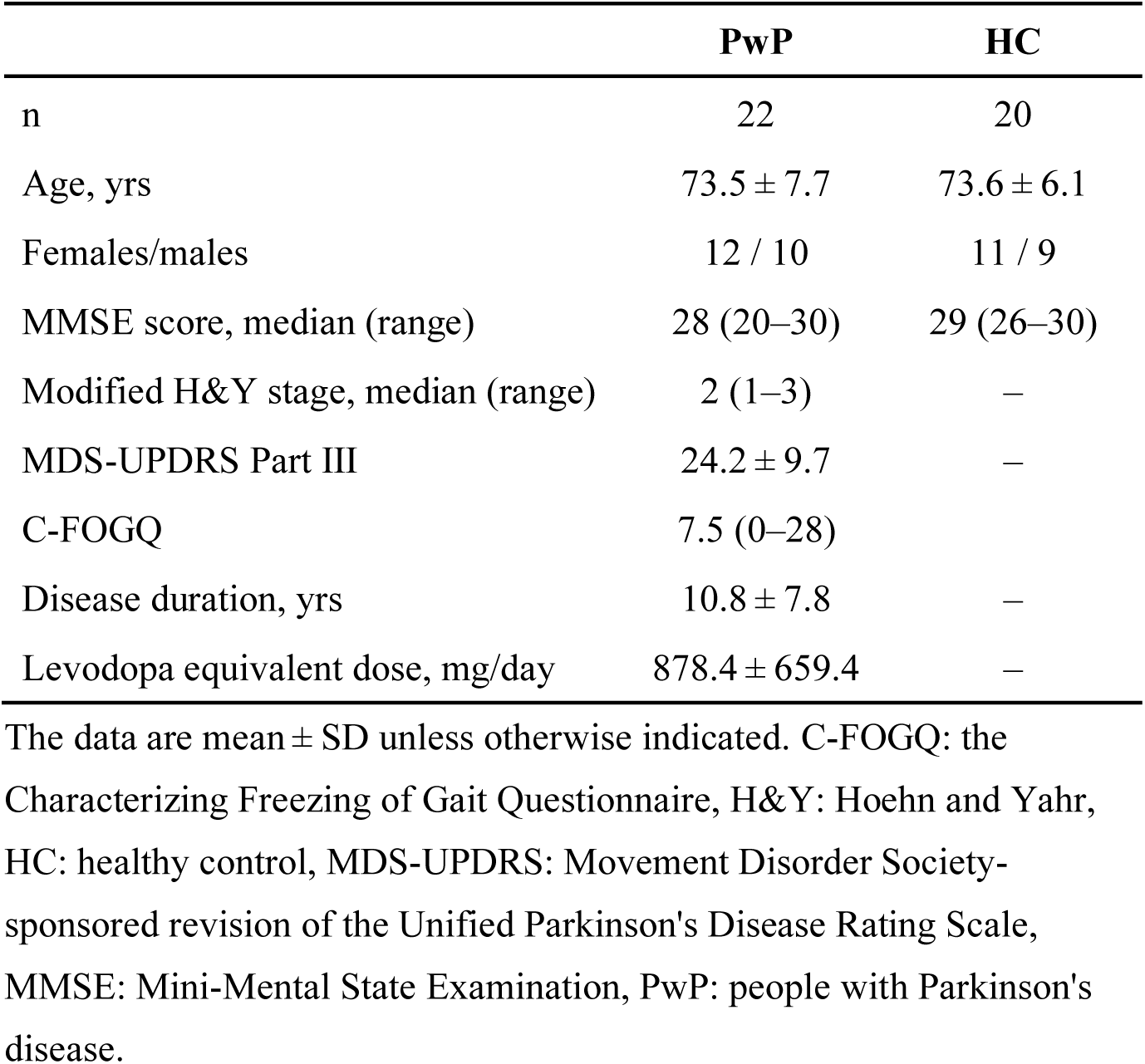
Demographic and characteristics of the HC and PwP.

|  | **PwP** | **HC** |
| --- | --- | --- |
| n | 22 | 20 |
| Age, yrs | 73.5 ± 7.7 | 73.6 ± 6.1 |
| Females/males | 12 / 10 | 11 / 9 |
| MMSE score, median (range) | 28 (20–30) | 29 (26–30) |
| Modified H&Y stage, median (range) | 2 (1–3) | – |
| MDS-UPDRS Part III | 24.2 ± 9.7 | – |
| C-FOGQ | 7.5 (0–28) | – |
| Disease duration, yrs | 10.8 ± 7.8 | – |
| Levodopa equivalent dose, mg/day | 878.4 ± 659.4 | – |

The data are mean ± SD unless otherwise indicated. C-FOGQ: the Characterizing Freezing of Gait Questionnaire, H&Y: Hoehn and Yahr, HC: healthy control, MDS-UPDRS: Movement Disorder Society-sponsored revision of the Unified Parkinson's Disease Rating Scale, MMSE: Mini-Mental State Examination, PwP: people with Parkinson's disease.

The inclusion criteria for HC were: (1) the ability to understand instructions, and (2) the absence of visual impairments or lower limb symptoms that could interfere with standing tasks. For the PwP, the inclusion criteria were: (1) a definitive diagnosis of PD according to established clinical criteria [24], (2) age ≥20 years, (3) if motor fluctuations (wearing off) were present, confirmation that the participant was in the ON state at the time of the study, and (4) the absence of visual impairments that could interfere with standing tasks.

The exclusion criteria were: (1) having a disease other than PwP that affects the central nervous system, (2) the presence of severe psychiatric symptoms such as hallucinations, (3) motor fluctuations not associated with medication timing, and (4) a sudden worsening of PD symptoms within 1 week prior to the study.

### Experimental procedure

For the experiment, the participants were seated on a chair or in a wheelchair and first completed a FOG questionnaire. They then stood on a force plate for the assessment of balance ability and then performed the standing CDT. The entire session lasted approximately 30 min.

### Control detection task

The CDT [21] was used to assess each participant’s motor awareness. This task has been employed to evaluate sensorimotor self-recognition in both healthy and clinical populations, including individuals with developmental coordination disorder [25,26]. The CDT was conducted using a personal computer with a 23.8-inch LCD monitor (1920×1080 pixels, 60 Hz; GW2470H, BenQ, Taiwan), a keyboard, and a force plate (TFG-4060, TechGihan Corp., Kyoto, Japan). Two black markers (a 6 mm diameter circle and a 6 mm square) were presented on the screen. When the participants moved their COP, both markers moved, but the relationship between the COP movement and marker trajectory (i.e., direction) varied across the trials.

The participants were instructed to shift their COP for a distance of up to 2 m and to judge which of the two on-screen markers they were controlling, as illustrated in Figure 1. Of the two markers, one continuously followed a prerecorded COP trajectory from another individual and was independent of the participant’s movement. The other marker moved according to a weighted combination of the participant’s real-time COP movement and the prerecorded trajectory, with the contribution of the participant’s own movement (control ratio, *c*) set at 20%, 40%, 60%, 80%, or 100% (100% indicating purely participant-driven motion). The prerecorded trajectory was extracted from COP data of a healthy individual recorded for 800 s prior to the experiment, with the starting position randomly selected for each trial. The position of the blended marker was updated based on the weighted sum of the velocity vectors of the COP and the prerecorded COP trajectory, using the following equation:

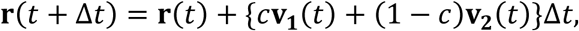

where **r**(*t*) ∈ ℝ²ˣ¹ denotes the position of the movement-controlled marker reflecting the participant’s motion, **v₁**(*t*) ∈ ℝ²ˣ¹ represents the velocity vector of the COP, **v₂**(*t*) ∈ ℝ²ˣ¹ represents the velocity vector of the prerecorded COP trajectory from another individual, *c* indicates the control ratio, and Δ*t* is the sampling interval.

**Fig. 1.**
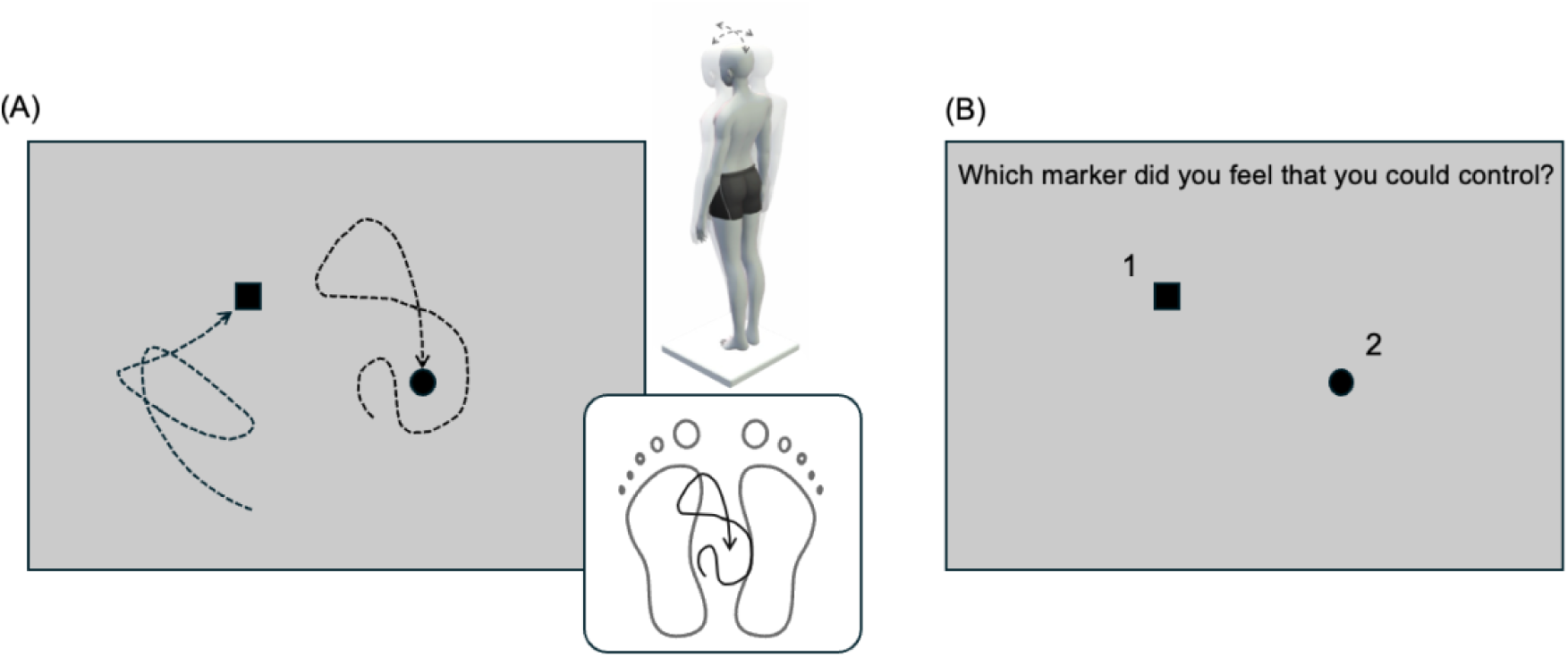
An example trial display for the control detection task. **(A)** Two markers (a *circle* and a *square*) moved on the screen in response to the participant’s center-of-pressure (COP) movement. **(B)** The participants judged which marker they were controlling based on the perceived correspondence between their voluntary COP movement and the marker trajectories.

If participants reached a confident decision before completing the 2 m COP displacement, they were allowed to respond and terminate the trial. When a trial ended — either after completing the 2 m movement or following a keypress — both markers stopped. A number (1 or 2) then appeared next to the markers along with the prompt, *“Which marker do you feel you could control?”* The participants responded verbally, and the experimenter recorded the response.

A total of 30 trials were administered in a randomized order, with each control ratio presented six times. Before the main task, the participants completed six practice trials (two each at 60%, 80%, and 100%) in random order. During the task, the COP trajectories were recorded continuously, and the total path length of each trajectory was calculated as an index of movement effort during the decision process. The response time was also recorded for each trial and defined as the interval between the trial’s onset and the space-bar press by the experimenter or when the participant completed the 2 m movement.

### Freezing of gait questionnaire

The severity of FOG in PwP was assessed using Section II of the Japanese version of the Characterizing Freezing of Gait Questionnaire (C-FOGQ) [27], which consists of 12 items addressing situations and frequencies in which FOG occurs. The participants responded to each item using a 5-point scale ranging from 0 to 4 (0 = not at all, 1 = rarely, 2 = sometimes, 3 = often, 4 = always), and the total score was used to quantify the severity of the participant’s FOG. Scoring of the C-FOGQ was conducted by a physical therapist based on an interview with each participant.

### Balance assessments

Static balance ability was assessed by recording COP trajectory while participants maintained quiet standing with eyes open on a force plate for 30 s. Postural stability during static standing was evaluated using the total COP path length and rectangular area. The rectangular area was calculated from maximum COP excursions in the left–right and anteroposterior directions.

Dynamic standing balance was evaluated using the Index of Postural Stability (IPS) (Figure 2) [28]. For this assessment, participants were instructed to shift their COP while standing to five target positions: forward, backward, left, right, and the center. COP data were recorded for 10 s at each position.

**Fig. 2.**
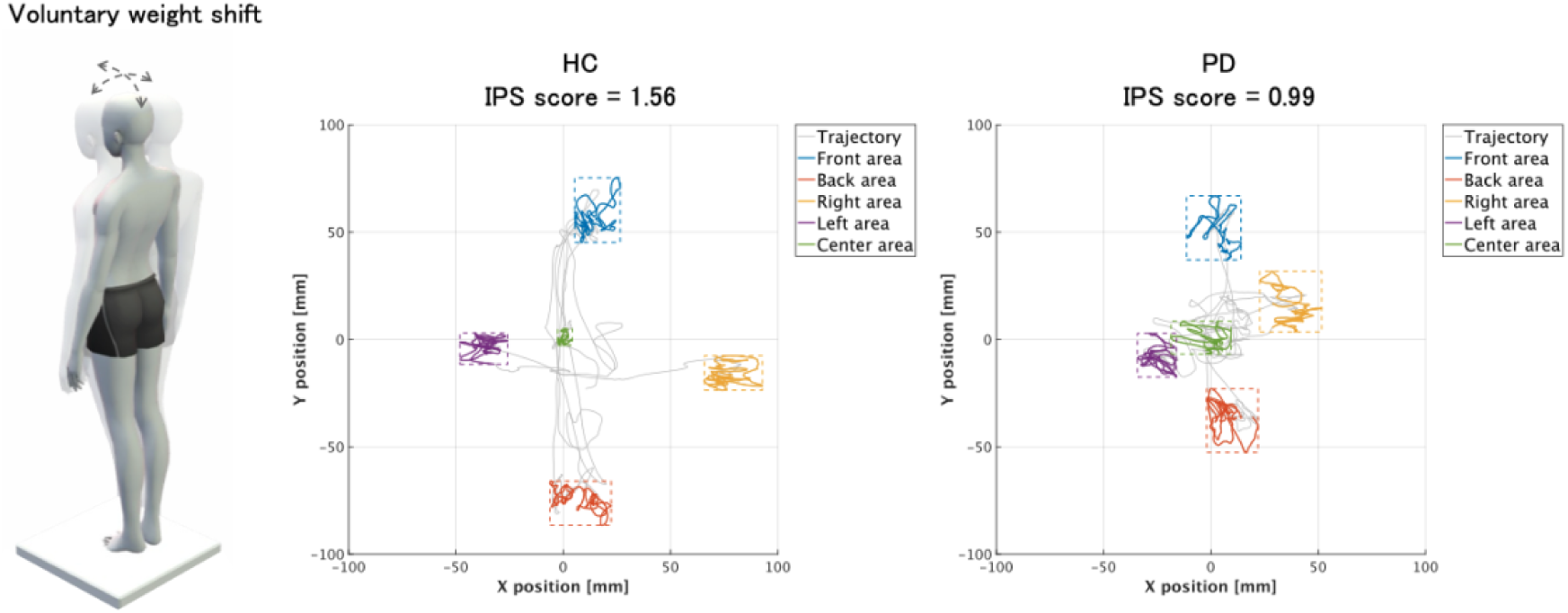
Representative COP trajectories during the Index of Postural Stability (IPS) assessment. The COP trajectories during voluntary weight shifting are shown for individuals in the HC (*left*) and PwP (*right*) groups. The participants were instructed to shift their COP from a central position to five target locations (front, back, right, left, and center) while standing on a force plate. Colored trajectories represent the COP sway recorded at each target position (front: *blue*, back: *orange*, right: *yellow*, left: *purple*, center: *green*), and *gray lines* indicate transitions. *Dashed rectangles:* the spatial region used to calculate the COP sway area at each position. The resulting IPS score is shown above each panel (HC: 1.56, PwP: 0.99), with lower values indicating reduced dynamic balance control.

The IPS results were calculated based on the average COP sway area at each position and the spatial extent of COP displacement across the directions, using the following equations:

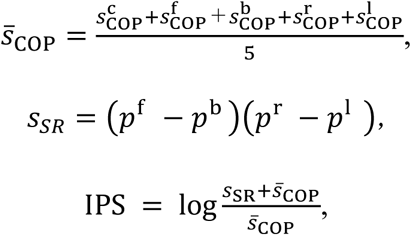

where *̅s*_COP_ represents the mean COP sway area across positions. The variables 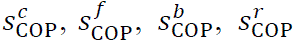 and 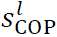 denote the COP sway areas measured during standing at the center position and during voluntary COP shifts to the forward, backward, rightward, and leftward positions, respectively. *s*_SR_ denotes the stability area defined by the distances between the COP positions in the anterior–posterior and left–right directions, and *p^f^*, *p^b^*, *p^r^,* and *p^l^* indicate the COP coordinates at the forward, backward, rightward, and leftward positions, respectively. Higher IPS values indicate better dynamic postural stability.

In all balance assessments, the COP data were recorded at a sampling frequency of 1,000 Hz using the above-described force plate. Prior to their analysis, the COP signals were low-pass filtered with a second-order Butterworth filter with a cutoff frequency of 20 Hz. Both static and dynamic standing measurements were performed twice, and the mean value across the two trials was used as the representative outcome.

### Clinical information

The clinical data of PwP were reviewed to obtain demographic and medical information. The collected data included age, sex, disease duration, severity of PD, levodopa equivalent daily dose (LEDD), and the results of neuropsychological assessments. Disease severity was evaluated based on the modified Hoehn and Yahr stage and the Movement Disorder Society-sponsored revision of the Unified Parkinson’s Disease Rating Scale (MDS-UPDRS) Part III. The neuropsychological assessments included the Mini Mental State Examination (MMSE).

### Statistical analyses

For the comparisons of the HC and PwP groups, independent-samples t-tests were conducted for variables that satisfied assumptions of normality. Effect sizes were estimated using Cohen’s *d*. To examine relationships between CDT performance and balance measures, Pearson’s correlation coefficients were used for analyses involving age and IPS, whereas Spearman’s rank correlation coefficients were used for static balance indices (i.e., COP trajectory length and rectangular area), which did not meet assumptions of normality. To further examine whether the association between CDT accuracy and balance performance could be accounted for by task-related movement extent during the CDT, partial correlation analyses were performed for the relationship between CDT accuracy and IPS while controlling for COP trajectory length during the CDT. In PwP, additional partial correlation analyses were conducted to examine whether the association between CDT accuracy and IPS was attenuated after controlling for clinical factors, including modified Hoehn and Yahr stage, MDS-UPDRS Part III score, and MMSE. To account for multiple comparisons, false discovery rate (FDR) correction was applied within each analytical set of balance measures.

To examine correlations between CDT performance and the clinical characteristics in PwP, we performed correlation analyses using Spearman’s rank correlation coefficients. This approach was used because several variables (e.g., the disease duration and LEDD) did not meet assumptions of normality; other clinical measures were ordinal in nature, including the MMSE, the modified Hoehn and Yahr stage, the MDS-UPDRS Part III score, and the C-FOGQ.

To confirm the adequacy of the sample size, we performed a post hoc power calculation using G*Power ver. 3.1. Based on the observed effect size for CDT accuracy (Cohen’s d = 1.10), the achieved power was 0.97, indicating sufficient sensitivity for HC–PwP comparisons. In addition, the post hoc power estimation for correlation analyses involving the CDT accuracy and static balance indices or the IPS score indicated adequate power (>0.80) to detect medium-to-large correlations.

The statistical analyses were performed using SPSS version 28 (IBM, Armonk, NY). The significance level was set at 5% for all analyses. The data are presented as the mean ± SD unless otherwise indicated.

## Results

### Accuracy, trajectory length, and response time on the CDT

The mean CDT accuracy across blending conditions was 79.8 ± 8.6% in HC and 65.3 ± 16.2% in PwP, a significant group difference (effect size Cohen’s *d* = 1.10, *p* < 0.01; Fig. 2A). The percentages of correct responses by control ratios in HC were 45.8% (self-control ratio 20%), 73.3% (40%), 85.8% (60%), 95.8% (80%), and 98.3% (100%), and those in PwP were 47.0%, 56.0%, 63.6%, 79.5%, and 80.3%, respectively.

The average COP trajectory length per trial, reflecting movement effort, was 1,213 ± 258 mm in HC and 922 ± 476 mm in PwP, a significant group difference (Cohen’s *d* = 0.75, *p* = 0.02; Fig. 2B). The trajectory lengths by control ratios in HC were 1,483 mm (self-control ratio 20%), 1,420 mm (40%), 1,213 mm (60%), 1,040 mm (80%), and 909 mm (100%), and those in PwP were 1059 mm, 982 mm, 920 mm, 858 mm, and 791 mm, respectively.

The average response time per trial was 8.5 ± 1.6 s in HC and 14.2 ± 6.6 s in PwP, a significant group difference (Cohen’s *d* = 1.3, *p* < 0.01; Fig. 2C). The response times by control ratio in HC were 9.5 s (self-control ratio 20%), 9.7 s (40%), 8.7 s (60%), 8.0 s (80%), and 6.6 s (100%), and those in PwP were 16.8 s, 14.4 s, 13.7 s, 13.6 s, and 12.6 s, respectively.

Correlation analyses revealed no significant relationships between the CDT accuracy and COP trajectory length, between the CDT accuracy and response time, or between the COP trajectory length and response time in either the HC, the PwP, or the combined cohort (all *p* > 0.05; Fig. 3A–C).

**Fig. 3.**
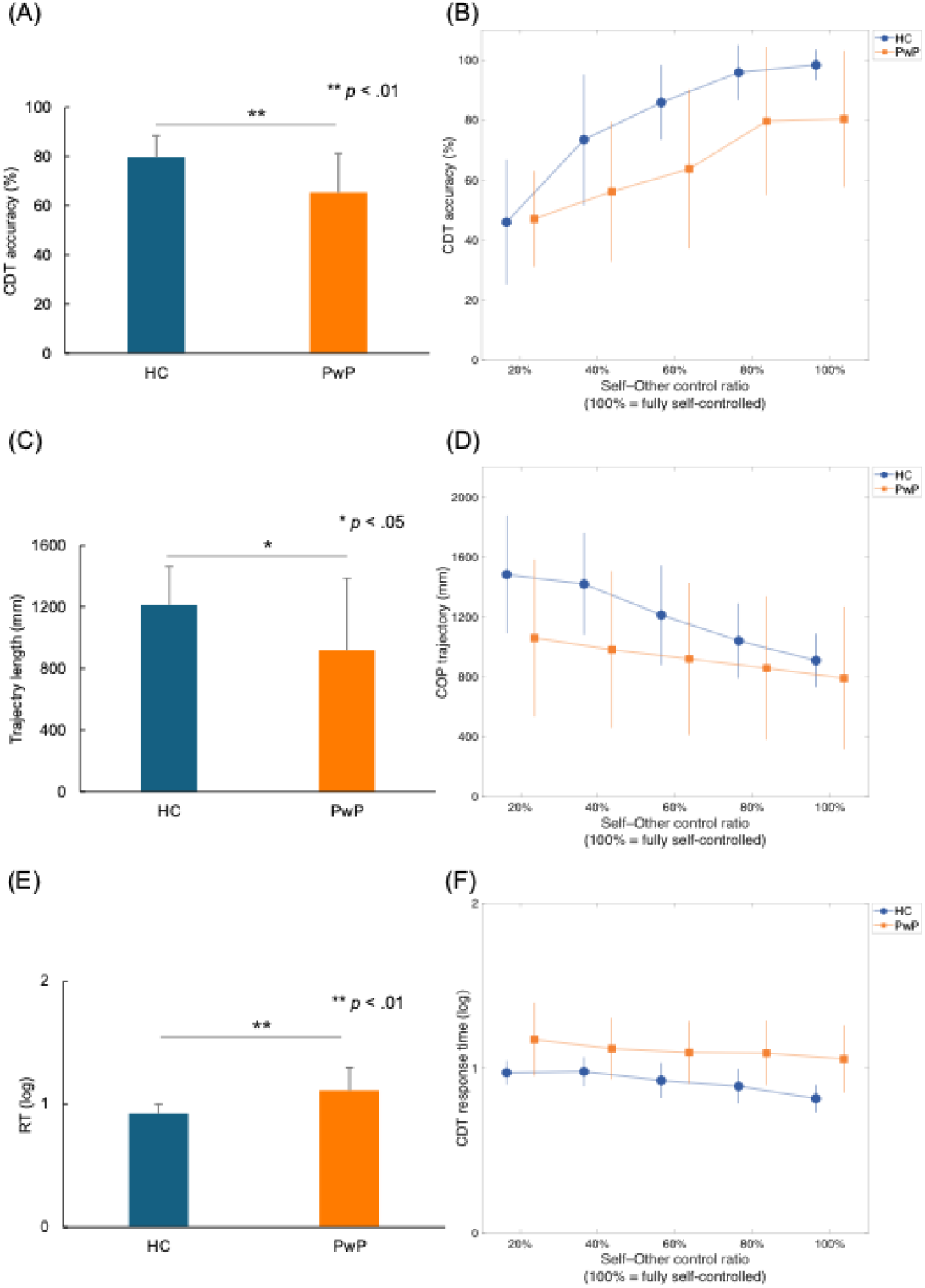
Group differences in the CDT performance and the effects of the self–other control ratio. **(A)** The mean CDT accuracy (%) across all control ratios for the HC and PwP groups. **(B)** The CDT accuracy as a function of the self–other control ratio (20%–100%, with 100% indicating fully self-controlled movement), for HC and PwP. **(C)** The mean center-of-pressure (COP) trajectory length during the CDT trials for HC and PwP, reflecting movement effort. **(D)** The COP trajectory length as a function of the self–other control ratio for each group. **(E)** The mean response time during the CDT trials for HC and PwP. **(F)** The response time as a function of the self–other control ratio for HC and PwP. Bars and points: group means. Error bars: standard deviation (SD). \**p* < 0.05, \*\**p* < 0.01.

**Fig. 4.**
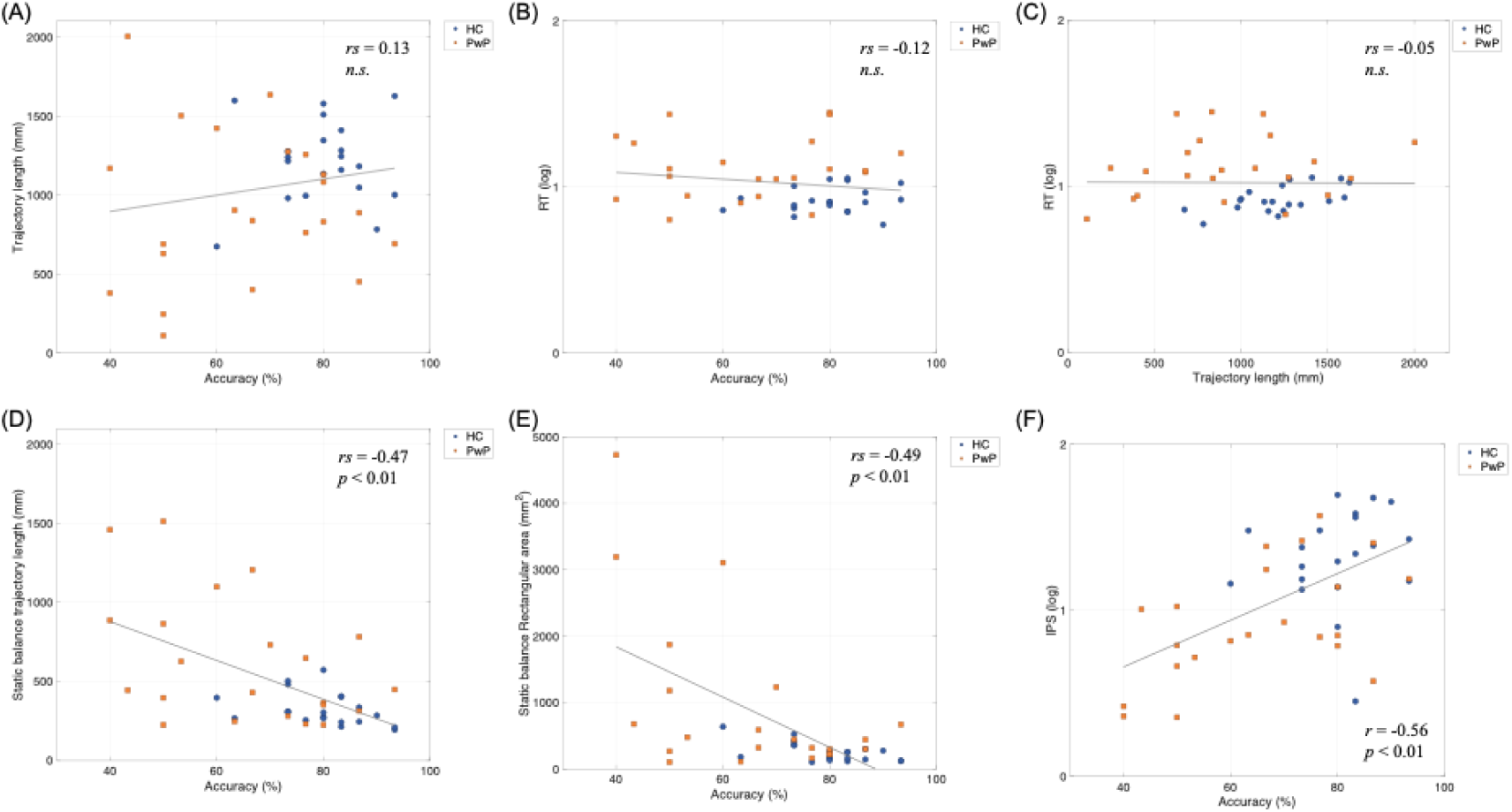
The relationships among the participants’ CDT accuracy, trajectory length, response time, and balance indices. Scatter plots illustrate the relationships between the CDT accuracy and the task-related measures or balance indices in the HC (*blue circles*) and PwP (*orange squares*) groups. The following relationships are depicted: **(A)** between the CDT accuracy and the COP trajectory length during the CDT, **(B)** between the CDT accuracy and response time, **(C)** between the COP trajectory length and response time, **(D)** between the CDT accuracy and static balance trajectory length, **(E)** between the CDT accuracy and static balance rectangular area, and **(F)** between the CDT accuracy and the IPS scores. *Solid lines:* regression lines for the combined cohort. Correlation coefficients and corresponding p-values are shown in each panel.

### Correlations between the CDT performance and balance indices

Correlation analyses examined the relationships between CDT accuracy and balance measures within the HC group, PwP group, and the combined cohort (Table 2, Fig. 3D–F). In the HC group, CDT accuracy was negatively correlated with both the trajectory length (*rs* = –0.48, *p* = 0.03) and the rectangular area (*rs* = –0.52, *p* = 0.02) in static balance, whereas the correlation with IPS scores was not significant (*r = –*0.14, *p* = 0.56). After FDR correction within the HC group, the relationships with trajectory length and rectangular area remained significant, and the relationship with the IPS score remained non-significant. A boxplot analysis identified one outlier with a markedly low IPS value in the HC group. After the exclusion of this participant, a Pearson’s correlation analysis revealed a small positive association between CDT accuracy and IPS (*r = –*0.28, *p* = 0.24), which did not reach statistical significance.

**Table 2.**
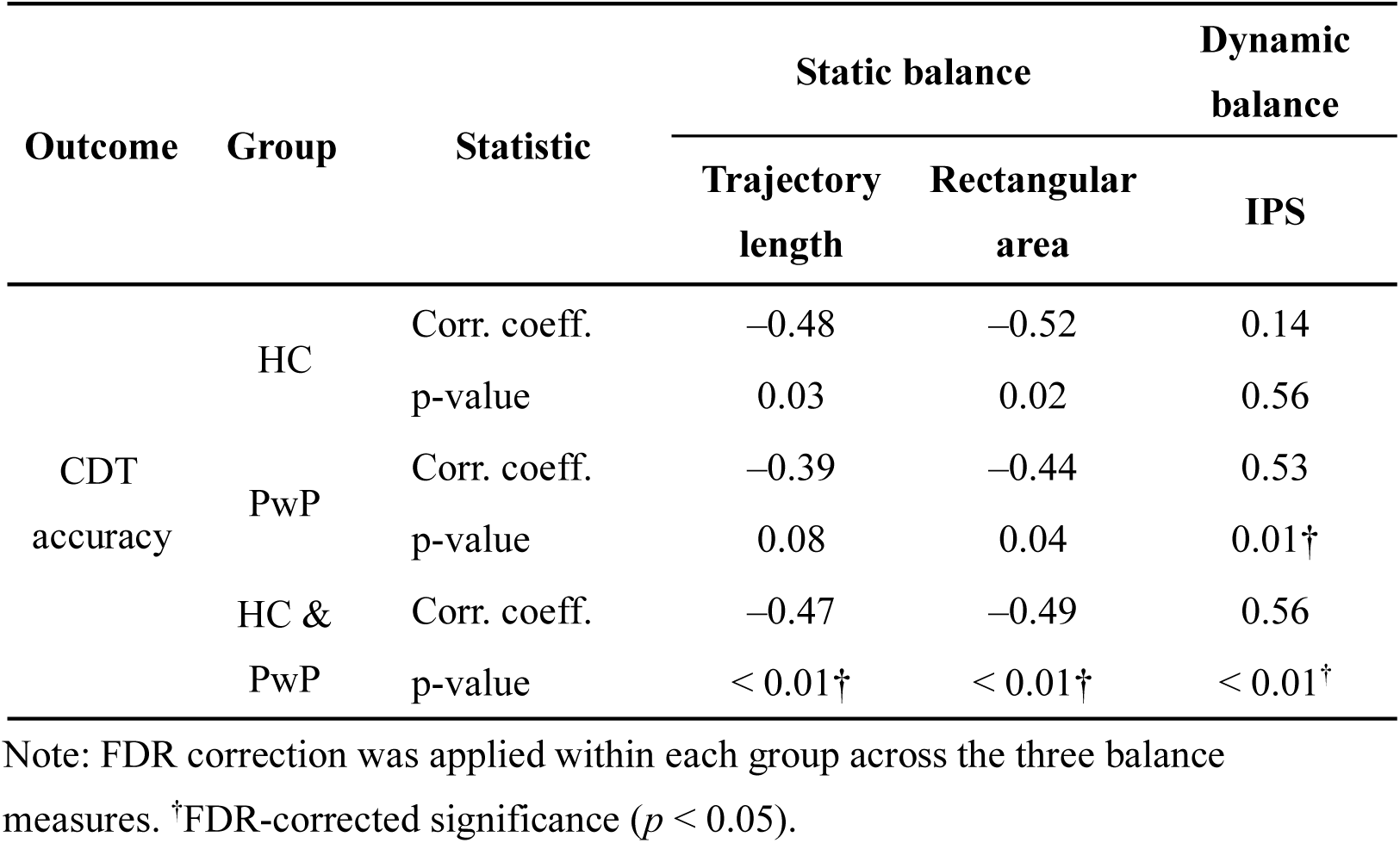
Correlation coefficients between CDT accuracy and balance measures.

In the PwP group, CDT accuracy showed correlations with trajectory length (*rs* = – 0.39, *p* = 0.08), rectangular area (*rs* = –0.44, *p* = 0.04), and IPS score (*r* = 0.53, *p* = 0.01). We conducted partial correlation analyses to adjust for potential confounders. When controlling for the PwP participants’ modified Hoehn and Yahr stage, the association between CDT accuracy and IPS score was attenuated but remained of moderate magnitude (partial *r =* 0.40, *p* = 0.07). With a separate adjustment for the MDS-UPDRS Part III and MMSE, the associations remained significant (partial *r* = 0.52, *p* = 0.02 and partial *r =* 0.45, *p* = 0.04, respectively). However, when the modified Hoehn and Yahr stage, MDS-UPDRS Part III, and MMSE were entered simultaneously as covariates, the association between CDT accuracy and IPS score was further attenuated and no longer significant (partial *r =* 0.29, *p* = 0.22).

In the combined HC and PwP cohort, CDT accuracy was correlated with trajectory length (*rs* = –0.47, *p* < 0.01), rectangular area (*rs* = –0.49, *p* < 0.01), and IPS score (*r =* 0.56, *p* < 0.01), and all three relationships remained significant after FDR correction. Moreover, the association between CDT accuracy and IPS remained significant after controlling for COP trajectory length during the CDT (partial *r* = 0.54, *p* = 0.01).

### Correlation between accuracy in the CDT and characteristics

Correlation analyses examined the relationships between CDT accuracy and the clinical measures in the PwP group. A significant negative correlation was observed between CDT accuracy and the modified Hoehn and Yahr stage (*rs* = –0.46, *p* = 0.03). In contrast, CDT accuracy was not significantly correlated with the age (*rs* = –0.04, *p* = 0.86), the disease duration (*rs* = –0.10, *p* = 0.65), MDS-UPDRS Part III score (*rs* = –0.13, *p* = 0.57), the C-FOGQ score (*rs* = –0.13, *p* = 0.56), the MMSE score (*rs* = 0.27, *p* = 0.23), or the LEDD (*rs* = –0.15, *p* = 0.49).

## Discussion

We investigated whether altered motor awareness—defined as the evaluation and attribution of self-generated postural actions—is associated with balance ability in PwP using the CDT to assess self–other attribution during balance control. Correlation analyses revealed that poorer CDT accuracy was associated with impaired balance performance, with no significant associations with motor symptoms, cognitive function, or freezing-related symptoms. Thus, the participants with poorer balance exhibited a greater reduction in the ability to distinguish self-from-other motion, which was independent of their motor and cognitive status. To our knowledge, this is the first study to apply the CDT to COP control in Parkinson’s disease, thereby providing a novel behavioral perspective on balance instability.

Compared to the healthy controls, the PwP participants showed significantly lower CDT accuracy, longer response times, and shorter COP trajectory length during the task. The prolonged response times may reflect additional time required for motor adjustment in the context of underlying impairments such as bradykinesia or reduced postural automaticity. These findings may also indicate increased processing demands when a person with PD distinguishes between self-generated and externally generated motor outcomes. Although COP trajectory length differed between groups, the absence of significant correlations among CDT accuracy, trajectory length, and response time suggests that task performance was not explained solely by greater movement extent or slower responding. In addition, the association between CDT accuracy and IPS remained significant after controlling for COP trajectory length measured during the CDT in the combined cohort. These findings indicate that the relationship between CDT performance and balance ability cannot be fully explained by task-related movement extent alone, although a contribution of motor capacity cannot be excluded. This interpretation is consistent with evidence concerning altered movement perception in PwP [20].

Beyond group differences, CDT accuracy was associated with balance measures. Static indices (i.e., the trajectory length and rectangular area) were correlated with CDT accuracy in the HC group and the combined cohort, whereas dynamic balance, indexed by the IPS, showed a significant association primarily in PwP. These findings suggest a link between balance performance and balance-related action–outcome monitoring, although the cross-sectional design precludes causal inference. Reduced balance stability may therefore reflect an altered evaluation of balance control in addition to motor execution deficits. In supplementary analyses, the association between CDT accuracy and the IPS was attenuated after adjustment for disease-related variables, and although it remained significant when adjusting separately for MDS-UPDRS Part III or MMSE, it was no longer significant when all covariates were entered simultaneously. The effect size, however, remained small to moderate, and the limited statistical power in the PwP group may have contributed to this attenuation. Overall, disease severity and cognitive status may partly explain the association, but a relationship between motor awareness and balance ability cannot be excluded. Larger studies are needed to clarify this relationship’s independence and underlying mechanisms.

Motor learning in rehabilitation relies on feedback-based error detection and the modification of motor commands, requiring an accurate awareness of discrepancies between intended and actual movement outcomes [29]. The association between CDT accuracy and balance measures suggests that altered motor awareness may constrain feedback-based learning in PwP. If balance errors are not accurately perceived or attributed, opportunities to adapt compensatory strategies may be reduced. Although causality cannot be established, we speculated that impaired motor awareness may limit the optimization of balance strategies through feedback-driven learning.

Our observation of dissociation between the static and dynamic balance measures provides additional insight. Static indices showed relatively consistent associations with CDT accuracy across the groups, whereas the IPS score was significantly correlated only in the PwP group. Because the IPS reflects effective use of the stability region during voluntary COP shifts, this pattern may indicate that motor awareness becomes more closely coupled with balance capacity under disease-related conditions that reduce compensatory redundancy. Alternatively, it may reflect balance impairment partly related to vestibular dysfunction in PwP [30], which compromises the reliability of sensory information that is critical for balance control. Under such conditions, dynamic balance may rely more heavily on the accurate prediction and monitoring of postural actions [31]. Thus, the IPS may capture both dynamic capacity and compensatory control in PwP, whereas in healthy individuals its contribution may be less pronounced under similar task demands.

Our analyses demonstrated that CDT accuracy was not significantly associated with conventional clinical measures, including the MDS-UPDRS Part III, MMSE, LEDD, or the C-FOGQ. Although it was related to disease stage (modified Hoehn and Yahr), the lack of associations with motor severity and global cognition suggests that CDT accuracy (*i*) captures aspects of motor awareness that are not fully reflected by standard clinical scales and (*ii*) may be more closely linked to functional balance capacity. We hypothesized that there would be an association with freezing of gait; however, no relationship with C-FOGQ scores was observed in this study. This may reflect limited sensitivity of the questionnaire-based measures and/or the relatively mild freezing severity in this cohort. Given the multiple reports linking freezing to sensorimotor integration deficits [16,32], it is clear that studies including a broader range of freezing severity are warranted.

From a clinical perspective, our findings suggest that altered motor awareness assessed by the CDT provides information that is not captured by conventional motor or cognitive measures in PwP. Because CDT accuracy was more closely related to functional balance measures than to standard clinical scales in the present study, we suspect that CDT accuracy may reflect how PwP monitor and evaluate their balance control during action. This distinction is relevant for rehabilitation, as postural instability often arises from impaired processing of sensory feedback and action outcomes rather than motor execution deficits alone [33,34]. Interventions emphasizing performance monitoring and structured feedback during balance tasks may therefore complement conventional training for PwP.

This study has several limitations. First, the PwP group showed a relatively restricted range of clinical severity, which may limit the statistical power to detect associations with specific clinical features. Second, although the CDT evaluates motor awareness via self–other attribution, specific components of the CDT were not systematically examined, and the influence of task-related motor demands cannot be fully excluded, even though the association between CDT accuracy and IPS remained significant after controlling for COP trajectory length measured during the CDT. In addition, sensory function was assumed to be largely preserved; however, more detailed assessments, including a peripheral neuropathy evaluation, are warranted in future studies to clarify the potential contribution of sensory factors. Finally, our study’s cross-sectional design precludes causal inference. Longitudinal and interventional studies are needed to clarify whether altered motor awareness contributes to balance impairment or reflects shared underlying mechanisms.

## Conclusion

Our findings demonstrate that altered motor awareness, assessed using the CDT, is associated with balance performance in PwP. Balance instability may therefore involve impaired motor awareness of balance control in addition to motor dysfunction, highlighting motor awareness as a complementary dimension for balance assessments and rehabilitation. These results support the exploration of motor-awareness-based approaches in balance rehabilitation, although their clinical relevance requires confirmation in further research.

## Acknowledgments

We thank the rehabilitation therapists at the Rehabilitation Center of Shiobara Hot Spring Hospital, Tochigi Medical Association for their assistance with the data collection.

## Statements and declarations

### Ethical considerations

This study was performed in line with the principles of the Declaration of Helsinki. Approval was granted by the Ethics Committee at The University of Tokyo (approval number: KE23-68).

### Consent to participate

Each participant provided written informed consent prior to participation.

### Consent for publication

Not applicable.

### Data availability statement

The data presented in this study are available from the corresponding author upon reasonable request, subject to institutional and ethical approval.

